# Effects of adhesive drying methods to reduce aerosol generation during resin bonding on enamel bond strength

**DOI:** 10.1101/2021.03.31.437826

**Authors:** Muhammet Kerim Ayar, Onder Yesil

**Author notes:** **Corresponding author:** Muhammet Kerim Ayar, Department of Restorative Dentistry, Faculty of Dentistry, Usak University, Usak, 64020, Turkey, and.

## Abstract

**Objective:** In order to reduce the amount of aerosol during the use of dental adhesives, which are widely used in minimally invasive procedures, the effects of air stream-free adhesive drying methods on the enamel bond strength of resin adhesive systems were evaluated.

**Materials and Methods:** The four adhesive drying techniques used were an air-stream, a micro-applicator, a cotton pellet and an absorbent paper. Adhesive systems were Single Bond Universal and Optibond All-in-one. The group in which the adhesive was not dried served as the negative control group. Enamel shear bond strength was performed with a universal tester with a crosshead speed of 1 mm/min (*n*=12). A two-way analysis of variance and the Tukey post-hoc test were used for analysis of the SBS data.

**Results:** For both adhesives, drying the adhesive with cotton pellet or micro-applicator provided a SBS mean values similar to air-stream drying, while statistically significantly lower SBS mean values were observed in the negative control group and in the absorbent paper-dry group compared to the air-drying group.

**Conclusions:** Drying the adhesive with micro-applicator and cotton pellets creates less aerosol and thus can be used in the COVID-19 pandemic as it provides enamel bonding strength similar to air drying.

**CLINICAL SIGNIFICANCE:** To provide safer dental care during COVID-19 pandemic, it is highly recommended to use non-aerosol-generating alternatives, instead of aerosol-generating dental procedures. In this study, we found that the aerosol generation-free methods for adhesive-drying such as cotton pellet and micro applicator drying methods provide similar enamel bonding with conventional drying method. We think that our findings will contribute to the realization of safer adhesive dentistry practice, especially during the COVID-19 pandemic.

## 1. INTRODUCTION

The COVID-19 disease caused by the Sars-Cov-2 virus was considered a pandemic disease by the World Health Organization (WHO) in early 2020.^1^ It is generally accepted that the COVID-19 virus is transmitted by contaminated surfaces, droplets and aerosols.^2^ Due to the extremely contagious nature of the virus, significant changes have been made in the provision of health care services in order to protect healthcare professionals and patients. Considering the transmission routes of the virus, it is recommended to limit the application of elective treatments in healthcare provision. Centers for Disease Control and Prevention (CDC) recommends the use of non-aerosol-generating alternatives to aerosol-generating equipment in dental practices, if any, in its guideline for the prevention of cross-infection for dentistry clinics in the COVID-19 pandemic.^3^

The provision of dental health care services with exception of dental emergencies, which were closed at the beginning of the COVID-19 pandemic, started to be reopened as a result of the reopening movement that took place worldwide in June and started to offer elective treatments in dentistry.^4^ However, it is generally accepted that as long as COVID-19, a pandemic transmitted by respiratory droplets, continues, it is not possible to continue dental services as before. Therefore, priority is given to researching new restorative dentistry procedures and methods that will minimize the generating of aerosol and droplets during emergency or elective dentistry treatments.^5^

During performing dental procedures, aerosols are released into the environment as a result of the use of some dental equipment including high and low speed rotary instruments, ultra-sonic scalers and air-water spray.^6,7^ If these aerosols released into the environment are contaminated with the virus, they may infect the dentist closest to the work area if he is not using the appropriate personal protective equipment correctly. These aerosols can also infect other people in the environment if the ventilation is not sufficient. For this reason, it is recommended to perform elective treatments with alternatives that do not generate aerosol instead of these aerosol-generating equipments.^6^

Performing of an adhesive restoration using dental adhesives requires the removal of the residual solvent step after the adhesive solution is applied to the tooth surface. Solvent removal is generally done by drying with oil-free compressed air stream. Since the air source is used during this process, the formation of aerosol carrying virus particles originating from the patient’s saliva may occur.^5^ For this reason, the techniques that will reduce the amount of aerosol that may occur during dental treatments using dental adhesive systems, especially during the COVID-19 pandemic, should be considered and the effect of these techniques on the quality of treatment should be evaluated.

The most common method of drying the solvent of the adhesive solution after it is applied to enamel or dentin surfaces is to dry with oil-free air stream.^8^ However, there are different adhesive drying techniques to assist this technique or as an alternative. Some of these are summarized in Table 1. Some of these methods are the prolonging the application time of the adhesive solution, the active application of the adhesive and the application of the adhesive with a warm air stream. Apart from these methods, even if they are not used as an alternative to air source after adhesive application until today, the adhesive solution can be dried with materials such as a cotton bellet, an absorbent paper and micro-applicator in order not to create aerosol during drying step of solvent. However, there is not enough information in the literature about how these drying methods, which do not use air stream during resin adhesive application processes, affect the bonding performance of adhesive systems. Therefore, the effect of non-aerosol-generating solvent drying techniques such as drying with cotton pellet, drying with absorbent paper and micro-applicator, on enamel bonding of two current universal adhesives was evaluated in the present study. The null hypothesis of the study is that “solvent drying methods have no effect on enamel bonding of the tested adhesive systems”.

**Table 1.** Studies about drying of dental adhesives on enamel bond strength.

| Reference | Adhesive system | Solvent drying technique(s) | Outcome |
| --- | --- | --- | --- |
| Tsuchiya et al. [12] | One-step self-etch adhesives | Adhesive application time (10 s, 20 s and 40 s) | The duration of the single-step self-etch adhesive application was a crucial factor for determining the enamel bond strengths of some adhesive systems tested. |
| Ikeda et al. [24] | One-step self-etch adhesives | Air-drying (0 s, 5 s and 10 s) | Air-drying was shown to have a significant effect on the removal of solvents included in the adhesive systems tested. |
| Shiratsuchi et al. [25] | One-step self-etch adhesives | Air-drying (normal vs warm) | Warm air-drying is essential to obtain adequate enamel bond strengths, although increasing the drying time did not significantly influence the bond strength. |
| Loguercio et al. [26] | Universal adhesives | Active vs passive application of adhesive system | Active application improved enamel bonding performances of universal adhesives when applied in self-etch mode. |
| Siqueira et al. [18] | Universal adhesives | Active application time (20 s vs 40 s) | Prolonged active application time of universal adhesive in self-etch mode increased enamel bond strengths. |

## 2. MATERIALS AND METHODS

### 2.1. Study design

The resin adhesive systems used in the present study were an universal adhesive system (3M ESPE Single Bond Universal, 3M Deutschland GmbH, Neuss, Germany) and an one-step self-etch adhesive system (Optibond All-in-one, Kerr, Kerr Italia S.r.I., Via Passanti, Italy). Enamel shear bond strength was a dependent variable. Drying resin adhesive after application with water-free oil-free air stream was deployed as a control group. Also, a group of no drying after application of resin adhesive served as a negative control group. The test groups of the study consist of the drying techniques of a resin adhesive with micro-applicator, cotton pellet or absorbent paper. Combinations of test groups, including control groups resulted in a total of 10 groups for the dependent variable (*n*=12).

### 2.2. Specimen preparation

Incisor teeth obtained from two-three-year-old bovines were used as a substitute for human teeth in this study. Teeth were kept frozen at −20 degrees for a maximum of 3 months before use.^9^ The vestibular surfaces of the crown of 120 bovine incisors were used after the root section and pulp removal. Teeth were randomly divided into ten groups according to the adhesive brand and adhesive drying techniques. The teeth were embedded in a hard plaster with a cylindrical mold, leaving the vestibule surfaces exposed, individually. Vestibule surfaces were first flattened with 240- grit silicon carbide (SiC) abrasive paper, then polished with 600-grit SiC abrasive paper for 60 seconds to obtain standard smear layers under water cooling. Debris on the polished enamel surfaces before adhesive processes was removed by cleaning with acetone-soaked cotton for 30 seconds.

A piece of double-sided tape with a 4-mm diameter hole was placed on the prepared enamel surface to limit the bonding area. According to the adhesive brand the prepared teeth were divided into two main groups. Then the teeth in each group were divided into 5 subgroups according to the drying technique. Adhesive systems were applied to the enamel surfaces in each subgroup according to the application procedures in the control and test groups shown below:

Subgroup 1: The adhesive systems were applied and dried by compressed air stream by air spray according to strict adherence to the manufacturer’s instructions (Table 2). These groups served as positive controls.

**Table 2.**
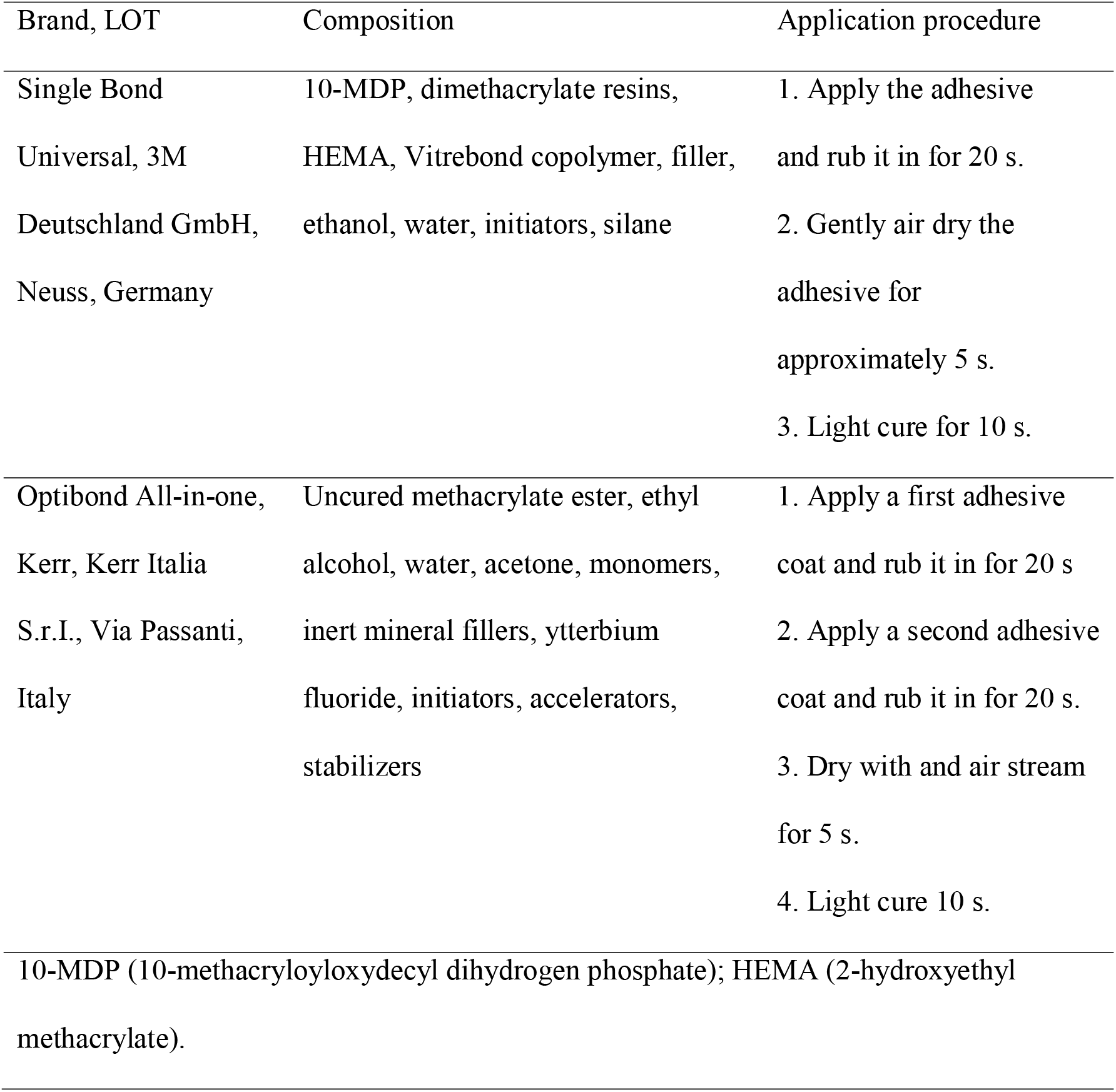
Materials used in the study.

Subgroup 2: Application of adhesive systems to enamel surfaces was done according to the manufacturer’s instructions. However, air drying was not done. These groups served as negative controls.

Subgroup 3: Adhesive systems were applied to the enamel surfaces according to the manufacturer’s instructions. However, the adhesive was dried by agitation with a new microapplicator for 20 s.

Subgroup 4: Adhesive systems were applied to the enamel surfaces according to the manufacturer’s instructions. However, the adhesive was dried with a piece of cotton pellet for 5 s.

Subgroup 5: As in the previous group, adhesive systems were applied to the enamel surfaces according to the manufacturer’s instructions, but the drying process of the adhesive was done by blotting with a piece of absorbent paper for 5 s.

The schematic design of study groups was also summarized in Figure 1. After the adhesive system application was completed, the resin composite buildups were prepared with a silicone mold with 2-mm diameter and 4-mm high on the enamel surfaces. The resin composite was placed into the mold in 2-mm thick layers and each layer was polymerized for 20 seconds with LED light curing device (1200 mW/cm^2^; Elipar S10; 3M Unitek, Monrovia, Calif).

**Figure 1.**
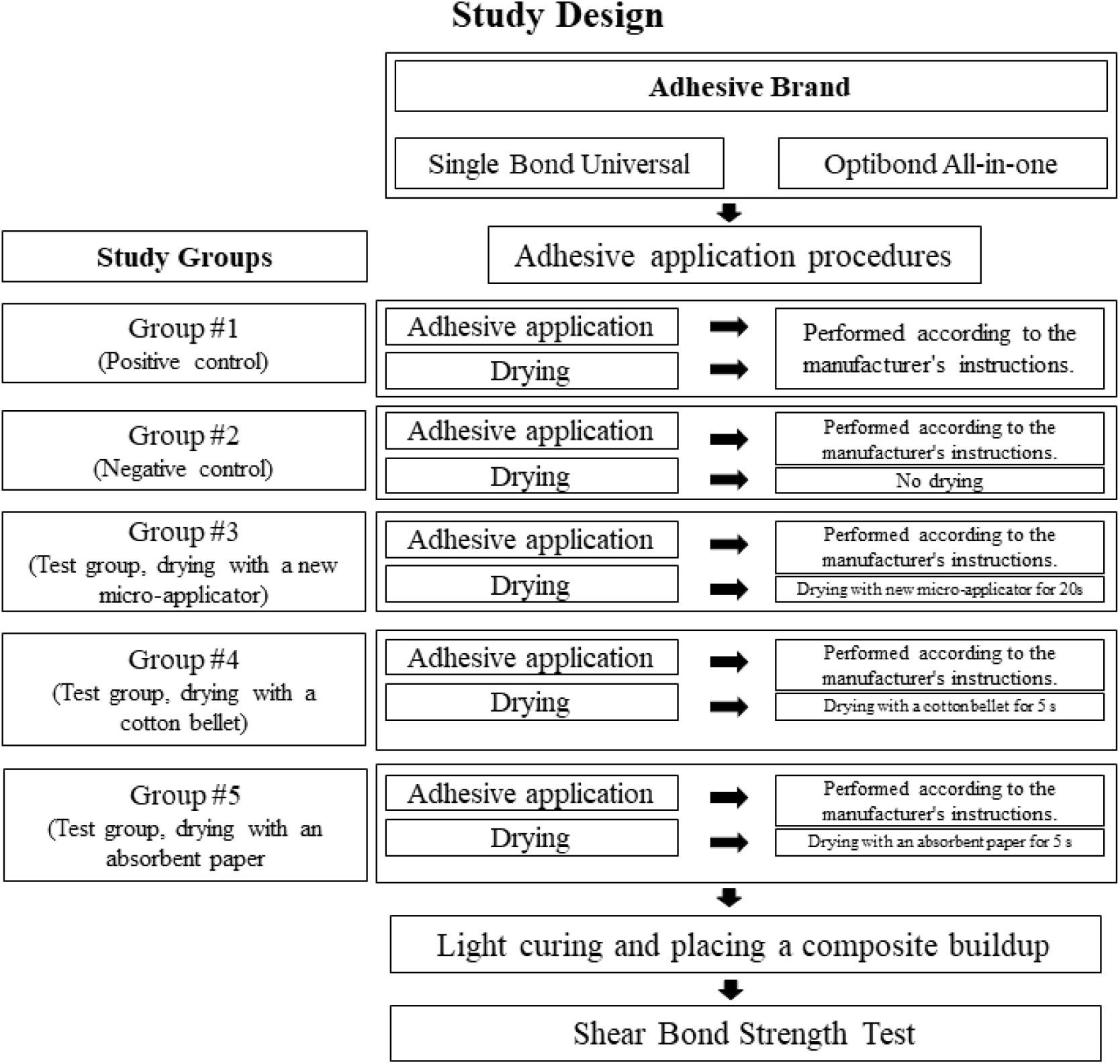
Schematic study design and groups of treatment.

### 2.3. Shear Bond Strength Test

The bonded samples were kept in distilled water for 24 hours prior to the bonding test. Shear bond strength test was performed with a knife-edge test apparatus on the universal test machine with a crosshead speed of 1.0 mm/minute with 12 samples for each group. Shear bond strength values were calculated in MPa by dividing the highest load value (N) recorded at failure the bonding area (mm^2^). After the bonding test, the failure mode was determined at 10x magnification with a stereo microscope (Meade Bresser Biolux, Meade Bresser, Rhede, Germany). The type of failure was classified as “adhesive failure” if the composite was not visible at all on the bonding surface, “cohesive failure” if it was in the composite or enamel, or “mixed failure” if it contained both structures.

### 2.4. Statistical Analysis

The results were analyzed by calculating the average shear bond strength (MPa) and standard deviations for each group. The difference between the shear bond strength averages was evaluated statistically by two-way analysis of variance (ANOVA; drying technique vs adhesive brand) and Tukey test at the 0.05 level of significance.

## 3. RESULTS

In Table 3, the shear bond strength (SBS) mean values and standard deviations, failure mode distributions for all adhesives and drying techniques are summarized. The highest SBS mean (24.53±3.6) was found in the positive control group of the Single Bond Universal adhesive system. The lowest SBS mean (13.22 ± 4.4) was seen in the negative control group of the Optibond all-in-one adhesive system. Two-way ANOVA revealed that there was no statistically significant interaction between the effects of drying technique and adhesive brand on SBS (*p*=0.971). The effects of both adhesive brand (*p*=0.00) and drying technique (*p*=0.00) on bond strength were statistically significant, respectively. For both adhesive systems, the positive control groups, in which the adhesives were applied according to the manufacturer’s instructions, showed the highest SBS mean values, and micro-applicator and cotton pellet drying techniques showed similar SBS mean values with the control group. For both adhesive systems, significantly lower SBS means were obtained in the negative control group in which the adhesive was not dried after application and in the absorbent paper drying technique group compared to the positive control groups. The frequency of different failure modes in SBS testing were shown in Table 3. The mixed failure mode was the predominant failure mode for all adhesives in positive control, cotton pellet group and micro-applicator groups, while adhesive failure was a predominant failure mode in in the negative control and absorbing paper groups.

**Table 3.**
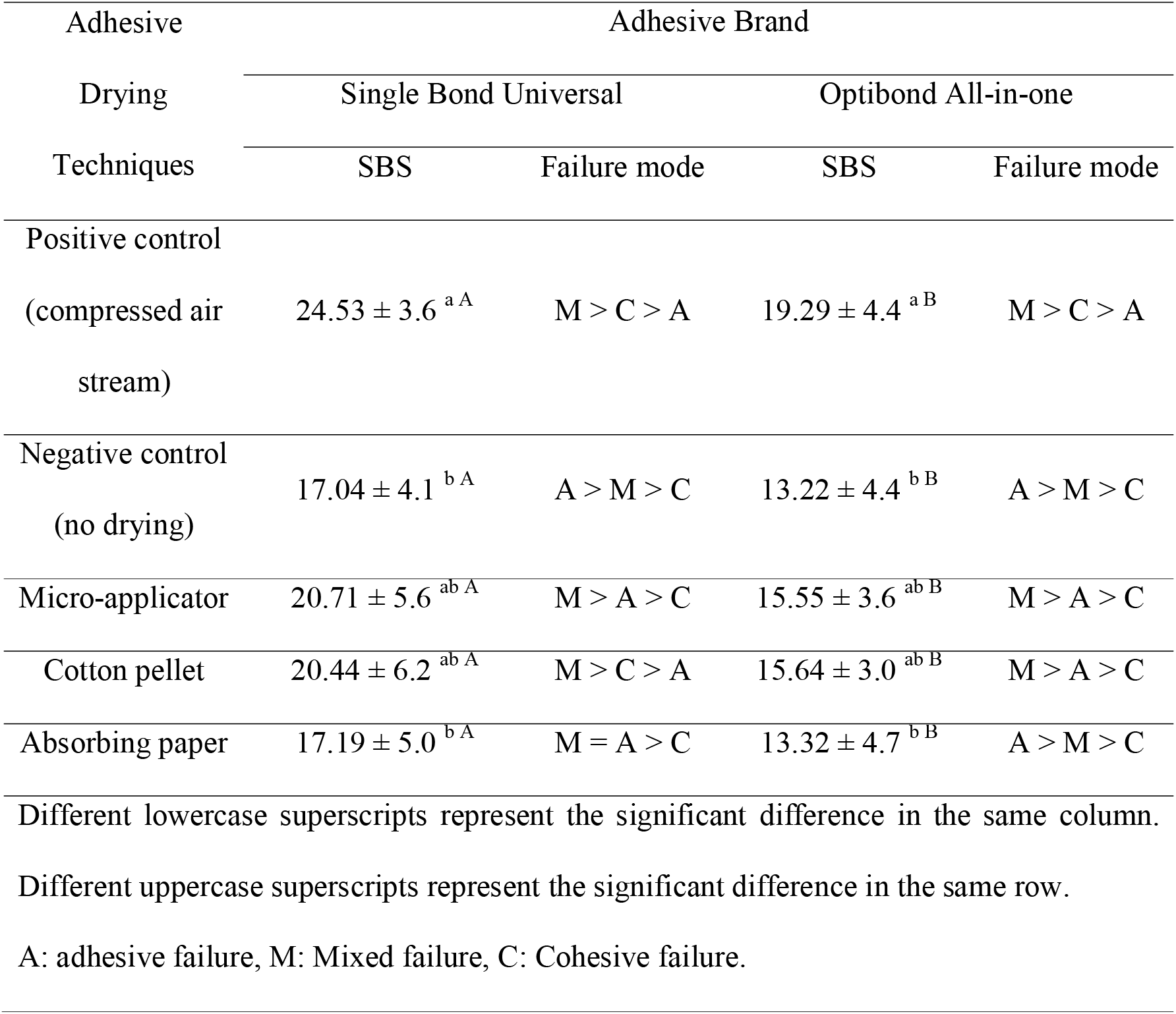
Shear bond strength (SBS) average and standard deviations and distribution of failure types (%) for all groups (*n*=12).

## 4. DISCUSSION

Considering the droplet and airborne ways of transmission of COVID-19 disease today, where the COVID-19 pandemic still continues, dental practices are at high risk for both dentists and patients in terms of the risk of covid-19 transmission.^6^ For this reason, national and international health authority organizations have published different guidelines that contain infection prevention measures specific to dentistry in order to minimize COVID-19 transmission in the dental environment and to protect staff. Among these preventive measures, in order to prevent the transmission of this infectious disease by aerosol, it is recommended to reduce the dental procedures that generate aerosol and to use alternatives to these procedures when necessary.^3^ In this context, in this study, the aerosol-free alternatives of drying the adhesive solvent with air spray, which is an operative procedure that creates aerosol during adhesive dentistry procedures, has been evaluated. The drying techniques included in the study were drying the adhesive for 20 seconds with a micro-applicator, drying the adhesive with cotton pellet for 5 seconds, and drying the adhesive with absorbent paper for 5 seconds.

The use of an adhesive resin system during placing the tooth-colored restorations ensures that the restoration will remain in the mouth for many years.^10^ Application of the adhesive system is a clinical procedure that requires more than one clinical step and any faulty clinical step will shorten the clinical life of the final restoration.^11^ One of these important clinical steps is to remove the residual solvents in the adhesive solution sufficiently after the adhesive solution is applied to the tooth surfaces. If there is much residual solvent left, the mechanical properties of the polymer created by the adhesive will be weak, and the bonding of the resin composite restoration with the dental tissues will not be strong enough.^12^ Therefore, the residual solvent of the adhesive should be removed according to the manufacturer’s instructions, which is ideal. According to the manufacturer’s instructions, this process is usually done with a compressed air stream by air spray.

In order for a technique to be an alternative to a conventional technique, the quality of the treatment performed with these techniques is expected to be similar to the quality of the treatment performed with the conventional technique. In adhesive applications, adhesive drying techniques that do not create aerosol are expected to provide bonding performance at a similar level to the enamel bond strength obtained by air spray drying technique, which is the conventional drying method. The main finding of the study is that the methods of drying the adhesive with cotton pellet and drying the adhesive with a micro-applicator provide similar enamel bonding strength with air spray drying.

There are some cases where cotton pellets are used during adhesive dentistry procedures. Moist cotton pellets are used to prevent overdrying of the dentin surface by air flow following acid-etching.^13,14^ Also, cotton pellet is used to keep the dentin surface moist before applying self-etch adhesives to the dentin following air-drying.^15,16^ Apart from that, there are also studies where the adhesive solution is applied with cotton pellets to the surface where it will be applied.^17^ However, the information in the literature about the use of cotton pellets to remove residual solvent of the adhesive is limited. The reason for this may be that dentistry has never encountered a situation where aerosol formation was extremely dangerous in dental practice, such as the COVID-19 pandemic. The dryness of the cotton pellet and the fibrous nature of the cotton may have enabled the adhesive applied surface to be dried effectively with cotton after the adhesive was applied to the enamel.

Another alternative that provides similar enamel bonding strength with air drying is to dry the adhesive by agitating it with a fresh micro-applicator for another 20 seconds. It has already been reported in the literature that prolonging the application time of self-etch adhesives to enamel increases the enamel bond strength.^12,18^ However, the application of this technique for drying the adhesive with a fresh micro-applicator was not available in the literature. Longer application of the adhesive may have increased solvent evaporation, and the use of a fresh micro-applicator may have provided the surface to be dried sufficiently by absorbing excess solvent by the dry micro-applicator.

Absorbent paper is also generally used in adhesive procedures to remove excess water from enamel and dentin surfaces.^19,20^ Especially when using etch-and-rinse adhesives, when the surface of the dentin surface should not be completely dried with air stream after acid-etching, therefore the dentin surface is dried with absorbent paper.^21^ In this study, absorbent paper was used to dry the remaining adhesive solution after the self-etch adhesive was applied to the enamel surface. According to the findings of the study, drying both adhesives with absorbent paper provided significantly lower enamel bond strengths when compared to airdrying control groups. This technique provided similar bond strength for both adhesives as in the negative control group in which the adhesive was not air dried after application. The reason for this finding may be that the absorbent paper would not absorb enough solvent from the surface. This finding suggests that drying with absorbent paper may not be appropriate way in order to minimize the amount of aerosol during adhesive procedures in clinical practice.

Enamel bond strength of two different adhesive systems was evaluated in the study. While one of these adhesives was a universal adhesive system and the other was a single-step self-etch adhesive, the universal adhesive was also applied in a self-etch mode. The reason for this was that pre-etching with phosphoric in the clinical setting during universal adhesive application would create a large amount of aerosol during acid washing process, which is against the recommendation of adhesive treatment by reducing the amount of aerosol.

In this study, the enamel bonding performance of adhesives applied in self-etch mode was evaluated. The application of universal adhesives and one-step self-etch adhesives to enamel without pre-etching with phosphoric acid results in lower bond strength, so enamel bonding of these adhesives is considered to be partially problematic.^22,23^ Therefore, it may be important to determine the effects of non-aerosol-forming adhesive drying techniques on enamel bonding, especially for self-etch adhesives.

